# VSEPRnet: Physical structure encoding of sequence-based biomolecules for functionality prediction: Case study with peptides

**DOI:** 10.1101/656033

**Authors:** Siddharth Rath, Jonathan Francis-Landau, Ximing Lu, Oliver Nakano-Baker, Jacob Rodriguez, Burak Berk Ustundag, Mehmet Sarikaya

**Author notes:** These authors contributed equally to this work (Jonathan Francis-Landau, Ximing Lu).

## Abstract

Predicting structure-dependent functionalities of biomolecules is crucial for accelerating a wide variety of applications in drug-screening, biosensing, disease-diagnosis, and therapy. Although the commonly used structural “fingerprints” work for biomolecules in traditional informatics implementations, they remain impractical in a wide range of machine learning approaches where the model is restricted to make data-driven decisions. Although peptides, proteins, and oligonucleotides have sequence-related propensities, representing them as sequences of letters, e.g., in bioinformatics studies, causes a loss of most of their structure-related functionalities. Biomolecules lacking sequence, such as polysaccharides, lipids, and their peptide conjugates, cannot be screened with models using the letter-based fingerprints. Here we introduce a new fingerprint derived from valence shell electron pair repulsion structures for small peptides that enables construction of structural feature-maps for a given biomolecule, regardless of the sequence or conformation. The feature-map introduced here uses a simple encoding derived from the molecular graph - atoms, bonds, distances, bond angles, etc., that make up each of the amino acids in the sequence, allowing a Residual Neural network model to take greater advantage of information in molecular structure. We make use of the short peptides binding to Major-Histocompatibility-Class-I protein alleles that are encoded in terms of their extended structures to predict allele-specific binding-affinities of test-peptides. Predictions are consistent, without appreciable loss in accuracy between models for different length sequences, marking an improvement over the current models. Biological processes are heterogeneous interactions, which justifies encoding all biomolecules universally in terms of structures and relating them to their functionality. The capabilities facilitated by the model expands the paradigm in establishing structure-function correlations among small molecules, short and longer sequences including large biomolecules, and genetic conjugates that may include polypeptides, polynucleotides, RNAs, lipids, peptidoglycans, peptido-lipids, and other biomolecules that could be implemented in a wide range of medical and nanobiotechnological applications in the future.

## Introduction

Cheminformatics tools have been used to predict solubility, binding-affinity to receptors, toxicity, and other properties of small-molecules, which, for example, include Extended-Connectivity-Fingerprints (ECFP’s) [1], Reduced-Graph representations [2], Simplified-Molecular-Input-Line-Entry-System (SMILES) [3], SMILES-Arbitrary-Target-Specification (SMARTS),[4] and International-Chemical-Identifier (InCHI) string analysis tools [5], Autoencoder implementations [6], Coulomb-matrices [7], Symmetry functions [8] and Graph-Convolutions [9,10]. Success of such tools have stimulated their implementation in bioinformatics. Graph-Convolution-Networks (GCN), where each amino-acid (AA) unit is considered as a node, has been used successfully on polypeptides as a classification tool in prediction of the protein-ligand interface [11]. Tools such as PotentialNet [12] that learn AA-connectivity of ligand binding sites have also been successfully implemented. The focus of such tools, however, has been on the small ligand and not the large biomolecular receptors. Additionally, a comprehensive structural feature-map is unavailable for proteins and peptides as neither the molecular structures nor their conformations are taken into consideration in the current GCNs. GCNs consider atom or AA connectivity for predicting properties of small-molecules. However, conformable biomolecules have connectivity beyond covalent bonds (such as hydrogen bonds) that are susceptible to changes based on the environmental and operational conditions. Tools directly employing three-dimensional coordinates as inputs to Neural Networks (NN) for small-molecule screening with integrated visualization-techniques have been developed [13]. However, the applications to biomacromolecules have been computationally intensive and currently impractical.

Traditional bioinformatics tools do not deal with small-molecules and are mostly concerned with AA sequences in proteins or oligonucleotide sequences in RNA and DNA. Letter-based representations are ubiquitous in addressing complicated functions owing to their simplicity, applicability, and accuracy in finding aligned domains in a sequence [14–17] or within a larger structure [18–20]. Several Machine Learning (ML) models to predict functionality using deep-learning, NNs, feature representation, and pattern analyses such as DeepMHC and NetMHCpan among others [21–23], have been developed by using the data in the Immuno-Epitope Database (IEDB) Analysis resource [24]. This database contains Major-Histocompatibility-Class-I, II (MHC-I, MHC-II) peptide-to-allele binding-affinity data for several species. In a recently developed Convolutional Neural Network (CNN), called DeepSeqPan [25], the authors recognize the importance of structural information in improving prediction accuracy and recommend their model as a supplement to other cumbersome models built with structural-alignment methods.

The traditional methodologies work only on letter-based AA or oligonucleotide labels and their derivations. The underlying physical-meaning, especially molecular structure or conformation is not apparent to the machine agent upon implementing ML algorithms. There is a loss of generalizability to include the molecules which do not have an obviously intrinsic sequence. Tools that work for or incorporate lipids, carbohydrates, and other biomacromolecules in their structures are exceedingly rare. Biological processes, however, are seldom isolated for a specific type of molecule, and commonly incorporate a wide range of biomolecules. Consequently, there is an imperative need for a method capable of encoding diverse biomolecules in a universal and meaningful manner (Fig 1) to study the interfacial phenomena at the molecular level. These processes may involve all biological systems, e.g., peptide and lipid or peptidoglycan [26], and biology/solid soft interfaces relevant to technological and nanomedicine applications [27].

**Fig 1.**
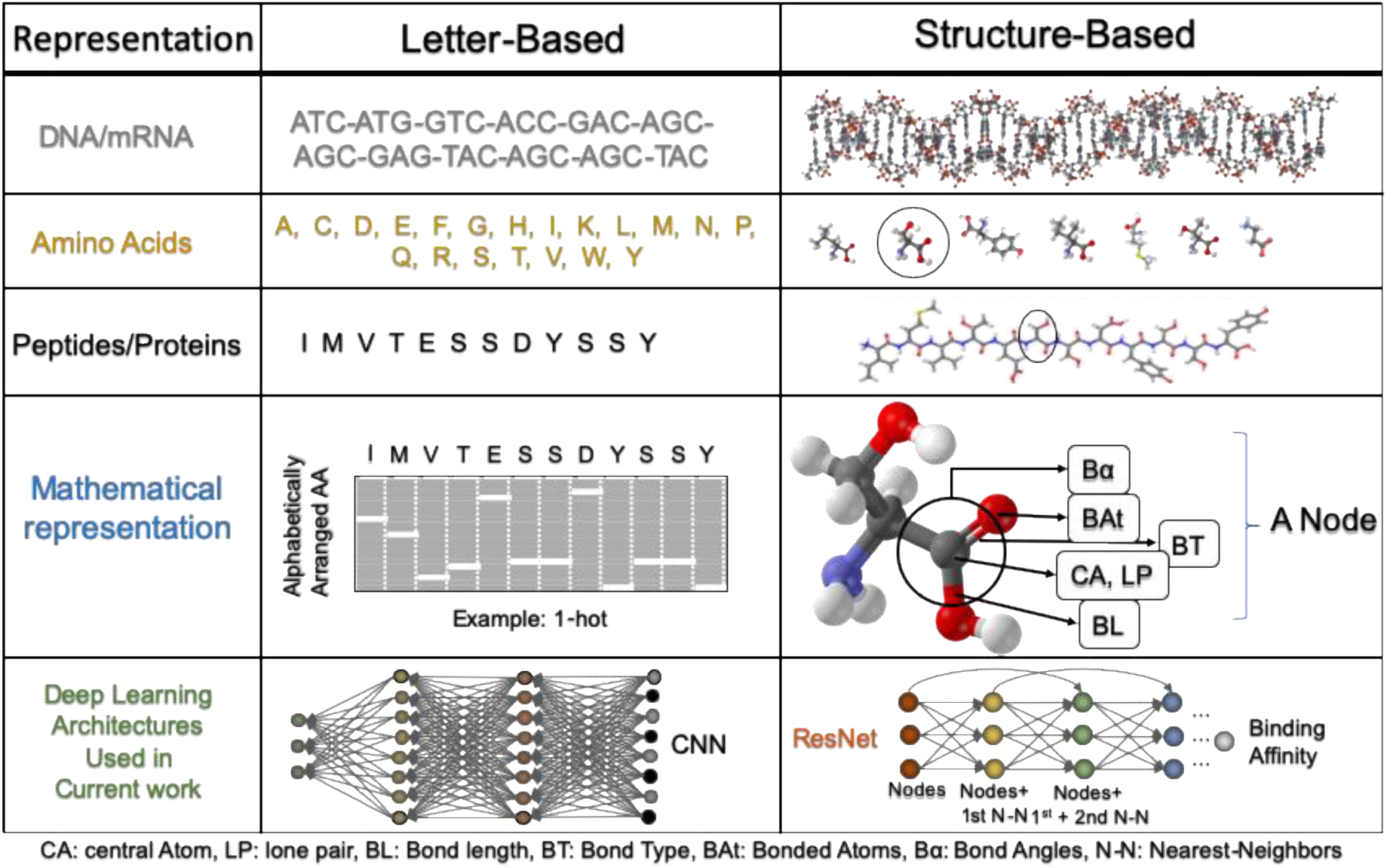
Schematics show the differences between the letter-based and structure-based representation of biomolecules for ML studies in functionality prediction. The central column is the index while the middle column shows the letter-based representation and the rightmost column shows the structure-based representation.

Implementations of such ML tools could broaden the paradigm of drug-design, combating antibiotic resistance, restorative dentistry [28], disease-diagnostics, biocompatible-coatings, lab-on-chip technologies, and biosensors [29]. In this work, we demonstrate a comprehensive feature-map for peptides that can be generalizable to other biomolecules. The immediate goals of the current work have been, (a) To take any AA sequence and convert it to a VSEPR structure-based representation via a reversible transformation; (b) To decide on an NN model that takes neighborhood information and performs consistently well across different length sequences, and (c) To benchmark the model with respect to the model used in DeepMHC. The long-term goal is to establish groundwork for future research in developing an accurate, interpretable and generalizable feature-map that incorporates conformations and multiple biomolecules to study complex phenomena.

The binding-affinity obtained from the current study displays higher prediction accuracy for 10-AA long peptides than the one-hot encoded shallow CNN model from DeepMHC [23], while the reverse is true for 9-AA long peptides. 5-fold Cross-Validation (CV) remains consistent across 9-AA and 10-AA long sequences, a significant improvement compared to DeepMHC where there is an appreciable drop in predictive power between 9-AA and 10-AA sequences. Since the VSEPR implementation consists of a larger feature map in conjunction with a deep residual neural network (ResNet), there is some overfitting and a loss of interpretability. It is noted that including angles in a GCN would be more interpretable. Indeed, such a model is aimed as one of the next steps to be taken towards generation of precise and pan-specific predictive tools, generalizable to other biomolecules of interest in medical and technological applications.

## Materials and Methods

### Data Cleaning and Preparation

Data compiled in 2013 from IEDB (www.iedb.org) Analysis Resource [24] are downloaded and cleaned. The binding affinities are measured in terms of Inhibitor Concentration IC50 required to reduce binding by half [30]. The values are converted to - ln(IC50), as a normalization step. According to extant standards, any sequence with an IC50 less than or equal to 500 is labeled as a binder and the others labeled as non-binder for binary classification. The dataset is then interfaced with a Python script to extract peptide sequences with transformed binding-affinity values to any allele of interest from any species within the dataset. All human alleles with at least 1,000 different corresponding epitope sequences are used in this study. 20% of the sequences are frozen out of the dataset for testing the model. Remaining 80% of sequences for each allele are used as a training set. The peptides in each set are then converted into their VSEPR encoded fingerprints as described below.

### VSEPR extended structure feature-map

As a first attempt, Bioluminate [31] is used to obtain the protein data bank (PDB) files for each of the naturally occurring AA. These PDB files contain information for each atom, including the data of atom type (in terms of atomic number) and cartesian coordinates of the given atom in space. The ProDy [32] library in python is used to traverse through the PDB files. Iterating through each of the neighbors of an atom, the bond type of each neighbor bonded to the central atom (CA) is obtained, based on prior knowledge of the AA structures. Euclidean distances are calculated to determine corresponding bond lengths. The number of lone pairs on any given CA is inferred based on the number of bonds and the bond-types that the given atom has, and its electronic structure. To calculate the angles made by pairs of Bonded-Atoms subtended at the CA, angle formula is used, and it is repeated parallelly for all CA and all combinations of Bonded-Atoms pairs per CA. Fig 2 shows the schematic of such a feature-map for Serine in an example peptide sequence.

**Fig 2.**
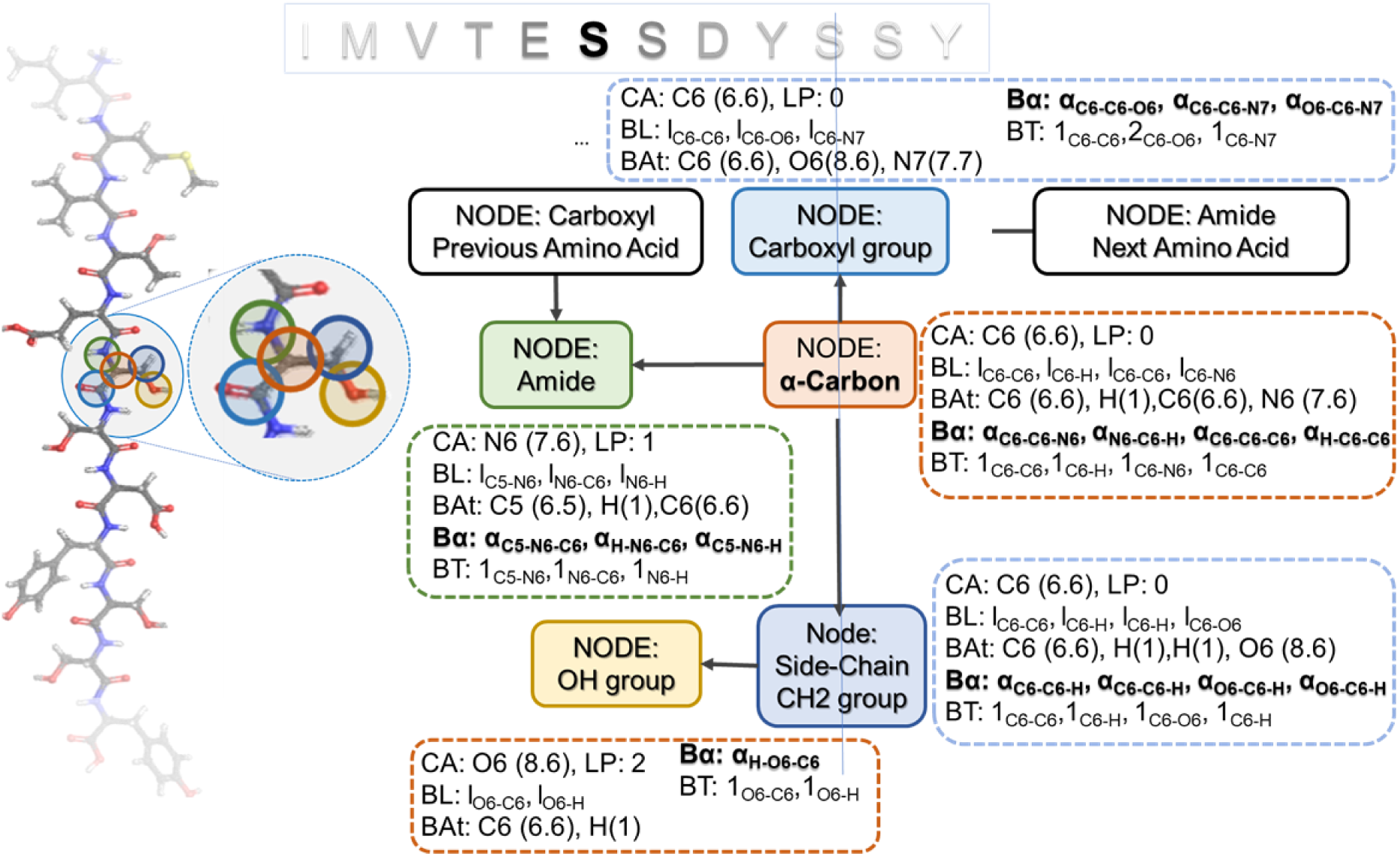
Schematic of Valence Shell Electron Pair Repulsion structural feature -map for bioinformatics studies. Green: N-terminus/Connection from previous Amino-Acid, Orange: Alpha-Carbon, Dark-Blue: Functional groups in side-chain, and Light-Blue: Connection to next Amino-Acid/C-terminus. Each such node contains 5 channels of information: Central Atom (CA) with associated Lone Pairs (LP), Bond lengths (BL), Bonded Atoms (BAt), Bond Types (BT) and Bond Angles (Bα).

The information for a given CA is appended to all successive non-hydrogen CA’s starting from the N-terminus of the peptide and ending at the C-terminus. Each type of parameter obtained from the VSEPR extended structure, is input as a separate channel of data to the neural network for training without overlap. The tenths place-value of the atomic number of the CA is the index of the residue location and the hundredths place-value is the location within the residue. For example, the α-Carbon in the 1^st^ AA at the N-terminus is given a value 6.01, whereas the carbon at the center of the planar carboxyl group bonded to the amine group of the 2^nd^ AA, is given a value of 6.00 (See S1 Appendix for more details).

The symbolic-connectivity reduces dimensionality but increases information bandwidth. It means that there are now two non-linear data bands in terms of power of 10. One band is 10^-2^ and the other is 10^-1^ in this case. Since the bands do not overlap, owing to the channel-splitting, machine learning methods also work as long as there are enough hidden layers to fit the respective non linearity levels. This is a way of multiplexing three separate inputs into one. Future implementations will eliminate this input through analytical transformation that only affects linear part of dominating input parameters. Binary vectorization of the encoding will also be attempted since power of two is more flexible instead of power of 10, in management of information bandwidth. Nevertheless, incorporating conformations as well as using adjacency matrices in a GCN is the clear next step towards making VSEPR methodology more impactful.

### Neural Network Architecture

Since behavior of molecular components of peptides depends on their neighborhood, Residual Neural Network (ResNet) was chosen to incorporate such information. The schematic of the process is shown in Fig 3. Such a Neural Network architecture comprises of a convolution block called the Residual Convolutional Unit (RCU) which performs a set of convolutions on the channels and a Fully Connected (FC) block. The RCU is implemented in terms of an Efficient Spatial Pyramid (ESP) [33]. ESP in the RCU allows for an improved gradient flow for training the network and essentially makes each atom ‘see’ its neighbors.

**Fig 3.**
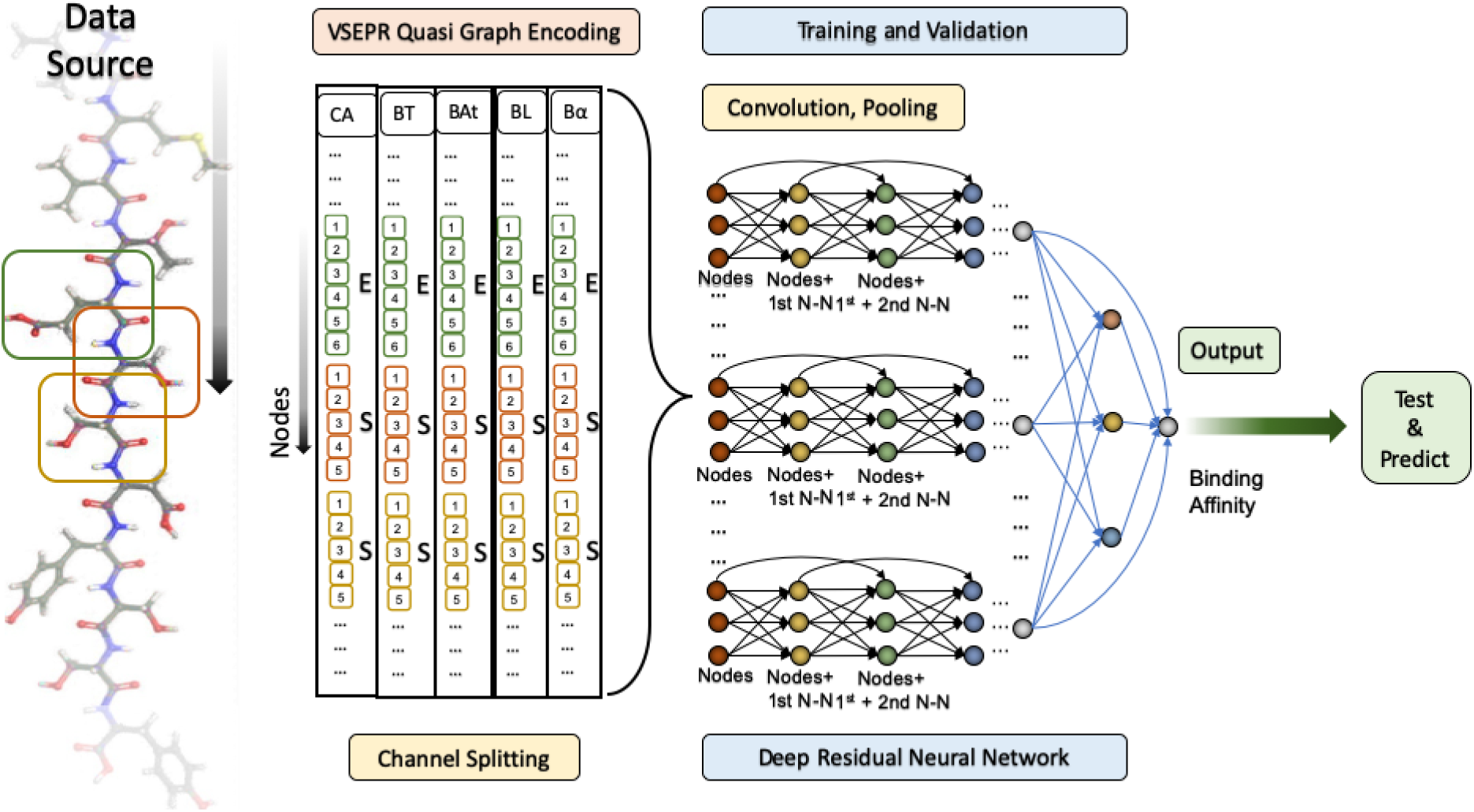
Schematic of the Training and Validation with the ResNet Architecture. In the convolution block, convolution proceeds on all atomic nodes simultaneously, with each successive layer seeing effects from more neighbors. Features thus extracted are sent through a fully connected network for prediction. The prediction can be carried out on any function that can be represented in terms of a numerical value. Here we predict the -ln(IC50) binding affinity.

The outputs of the ESP enhanced RCU block are then passed into the FC block, with a Rectified-Linear-Unit activation on all the layers and SoftMax on the last. Mean-Squared-Error is the loss function to be minimized to output the binding affinity of the peptide to the corresponding MHC-I allele. Batch Normalization is performed after every layer in the network. Randomly initialized weights are then learned in a supervised learning protocol and hyperparameters are tuned following a training process as described below.

### Training, Validation and Testing

Sequences for each allele in the training-set are divided into five equal parts randomly selected, to set the stage for a 5-fold cross-validation as a control against sampling bias. Four out of five such parts are used to train the model and the fifth one is used for testing. Then the model rotates through another set of four such parts as training and fifth one as test set. In each such model training round, per allele, each of the feature-maps are split into 5 channels per input sequence. They are sent in simultaneously in mini-batches of 20 peptides at a time into the ResNet described above, for 5000 epochs. The PyTorch [34] deep-learning library is used for training. The model is labeled ‘converged’, if validation loss (10% of the training data is used for validation) did not reduce by more than 1% for 100 subsequent epochs.

After the training is completed, hyperparameters are tuned to maximize the 5-fold cross validation resulting in a learning rate of 5e^-4^. The process is repeated three times to ensure that the cross-validations observed are consistent and not affected by choice of training samples. A similar procedure is followed to train a regular Convolutional neural Network with one-hot encoded peptide sequences as in DeepMHC for one-to-one comparison and evaluation. Meanwhile, the 20% of data frozen before training is then used as a blind test set for evaluating model performance.

## Results and Discussions

We compare the allele-specific VSEPRnet model with the state-of-the-art CNN model, DeepMHC that works with letter-based fingerprints. The 5-fold CV results as obtained by the reproduced DeepMHC model versus the current VSEPRnet model is shown in Fig 4A and 4B for 9-AA and 10-AA long sequences respectively. Results show a consistent response across sequence lengths in the VSEPRnet case in contrast to DeepMHC, where there is a fall in prediction accuracy for 10-AA sequences (refer S1 Fig). In the case of DeepMHC, the average 5-fold CV (Fig 4C) across all alleles studied is 0.87 for 9-AA sequences, with a standard-deviation of 0.03. For 10-AA sequences it is 0.65 with a standard deviation of 0.11. For VSEPRnet, the average 5-fold CV for 9-AA long peptides is 0.74 with a standard deviation of 0.06. While for 10-AA long peptides it is 0.69 with a standard-deviation of 0.03. Taking available data and overfitting into consideration, VSEPRnet therefore has a consistency in predictability over sequence lengths. One of the reasons for a marked fall in cross validation for 10-length sequences, as outlined in DeepMHC, is a dependency of the model on distal effects which dominate as lengths increase. We note that because feature-maps and neural-network architectures usually go hand-in-hand, further investigation is mandated to isolate the cause of the flattening response observed in the case of VSEPRnet. However, due to the nature of the ESP convolution block in the ResNet architecture, distal effects in the convolution may not dominate. Moreover, the distinction in input sizes between 9-AA and 10-AA peptides is based on physical rather than sequence length.

**Fig 4.**
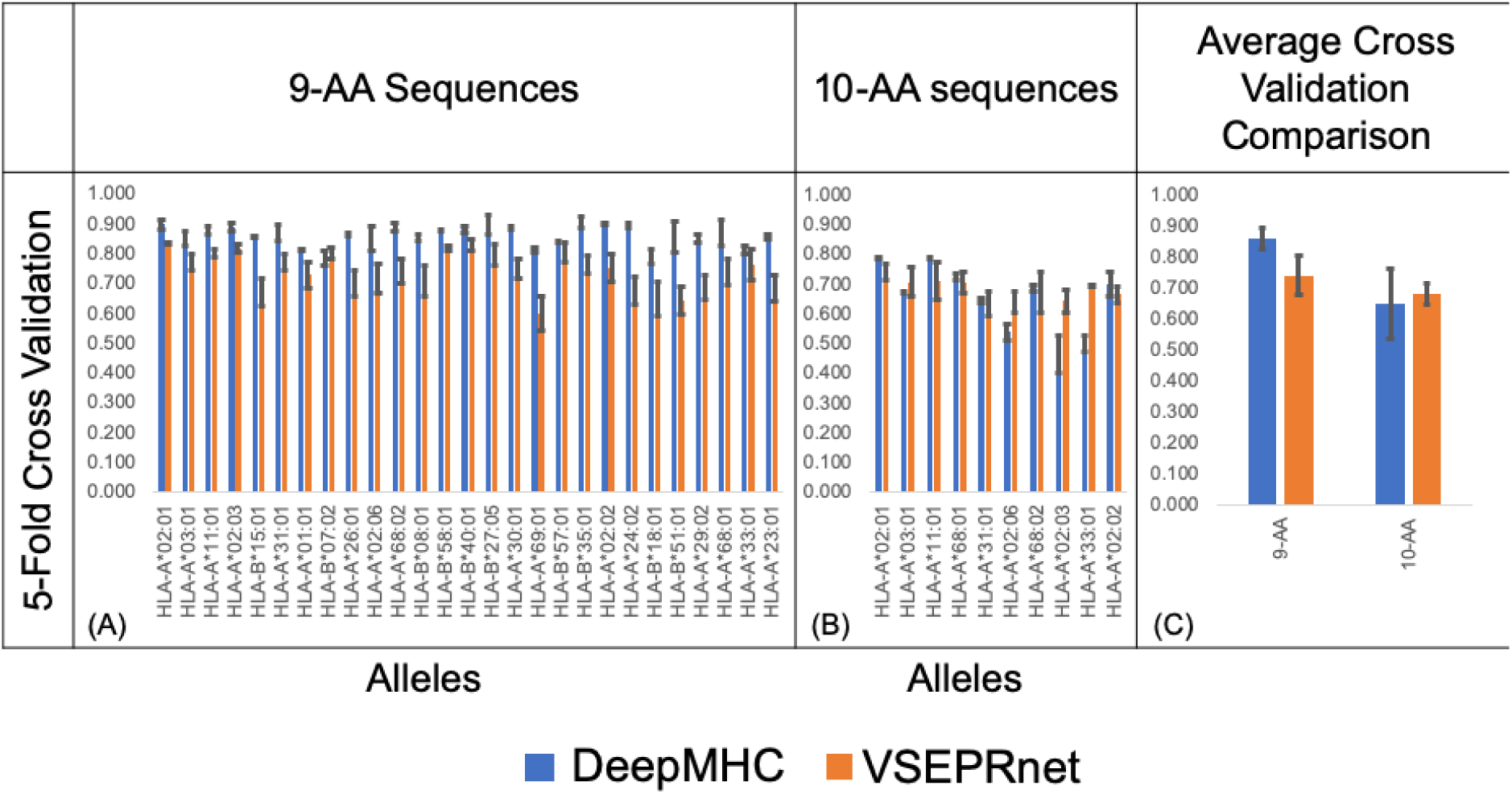
5-Fold Cross Validation results from VSEPRnet compared with DeepMHC. (A) 5-fold CV for 9-Length peptide sequences and (B) 5-fold CV for 10-Length peptide sequences; (C) Average 5-fold CV for 9 and 10-AA peptides across alleles. Performance of VSEPRnet falls in comparison to DeepMHC in the 9-Length peptides case for most alleles, the most probable reason being overfitting due to increased dimensionality. The performance of VSEPRnet is better than DeepMHC in case of 10-Length peptides on mostalleles due to reduced dominance of distal effects. Overall, VSEPRnet performs consistently across sequence lengths and does not have the drop in accuracy between 9 and 10-AA peptides as is the case with DeepMHC.

Since the VSEPR feature-map contains more information than the one hot encoding, the data required to avoid over-fitting becomes higher. Thus, the lack of requisite data-density lowers the average 5-fold CV from 0.87 (DeepMHC) to 0.74 (VSEPRnet) for the 9-AA long peptides. As discussed previously, there is a role-reversal for the 10-AA case because there is a pronounced distal-effect in the DeepMHC implementation whereas it is negligible for the VSEPRnet implementation (see S2 Appendix for more details). The overall performance of VSEPRnet in terms of 5-fold CV is contingent mostly on the available data-points to train on. Future work could be directed to implement the model on datasets with higher density of data obtained from High Throughput Sequencing techniques [35].

Performance Comparison of VSEPRnet and DeepMHC on previously frozen test data, uses Pearson Coefficient (PC), Spearman Rank Correlation Coefficient (SRCC) and Area Under receiver operating Curve (AUC) as metrics to compare the performance of the two models. For the PC metric, VSEPRnet wins on 11 out of 27 alleles in the 9-AA case and 5 out of 10 alleles in the 10-AA case; for the SRCC metric, VSEPRnet wins on 8 out of 27 alleles in the 9-AA case and 4 out of 10 alleles in the 10-AA case; And for the AUC metric, VSEPRnet wins or performs equally on 7 out of 27 alleles in the 9-AA case and wins on 9 out of 10 alleles in the 10-AA case. Across all metrics, the performance of VSEPRnet is within the first standard deviation of DeepMHC for 9-AA peptides, and for 10-AA peptides, VSEPRnet wins on both PC and AUC metrics. The average PC across all alleles for DeepMHC is 0.244 with a standard deviation of 0.112 for 9-AA peptides, and 0.279 with a standard deviation of 0.112 for 10-AA peptides. The average PC for VSEPRnet is 0.235 with a standard deviation of 0.096 for 9-AA peptides and 0.296 with a standard deviation of 0.042 for 10-AA peptides. Similarly, across all tested alleles, the average SRCC of DeepMHC is 0.6 with a standard deviation of 0.140 for 9-AA peptides and 0.514 with a standard deviation of 0.165 for 10-AA peptides, while the average SRCC, across all tested alleles for VSEPRnet is 0.571 with a standard deviation of 0.099 for 9-AA peptides and 0.5 with a standard deviation of 0.046 for 10-AA peptides (See Fig 5 and S2 Fig for more details).

**Fig 5.**
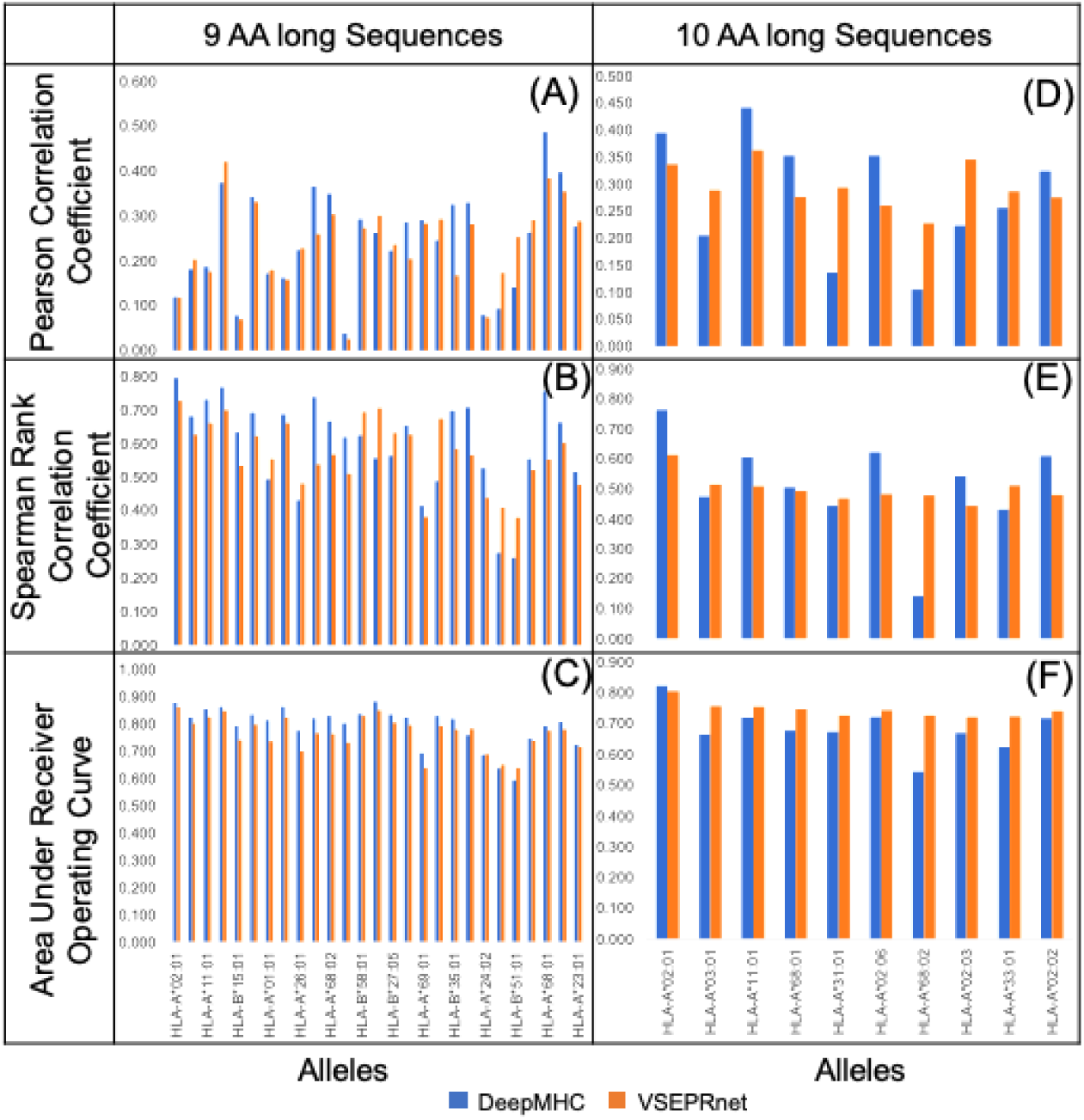
Comparison of DeepMHC and VSEPRnet on test data for 9-AA and 10-AA long peptides. DeepMHC performs consistently better on the (A) PC, (B) SRCC and (C) AUC metric for 9-AA long peptides because of lower probability of overfitting due to low information content of the 1-hot encoding. VSEPRnet PC values are within the mean and spread of the PC values for DeepMHC. For 10-AA, VSEPRnet performs equally as well as DeepMHC in case of (D) PC and (E) SRCC, while performing consistently better for (F) AUC owing to the elimination of distal effects.

Additionally, across all tested alleles, the average AUC of DeepMHC is 0.795 with a standard deviation of 0.072 for 9-AA peptides and 0.684 with a standard deviation of 0.072 for 10-AA peptides, while the average AUC, across all tested alleles for VSEPRnet is 0.767 with a standard deviation of 0.062 for 9-AA peptides and 0.745 with a standard deviation of 0.025 for 10-AA peptides. A consistent response across alleles is also shown by the VSEPRnet, without being affected by the sequence length of the peptides.

## Conclusions and Future Work

VSEPRnet is an introductory implementation for extending cheminformatics style feature-maps to bioinformatics studies while maintaining generalizability across lengths and molecule-types. There is a demonstrated consistency in prediction-accuracy of VSEPRnet model across alleles and between 9-AA to 10-AA long peptides binding to MHC-I allele. Therefore, there are advantages of using this implementation as a first step in generalization of feature-maps to include other molecules. There is a need to incorporate conformations and substrate information into the model to make it truly generalizable to DNA, RNA, proteins, peptides, intrinsically-disordered regions, lipids, peptidoglycans, phospholipids, sugars, and smaller biomolecules such as vitamins and co-factors.

Since the VSEPRnet 5-fold CV does not show appreciable dependency on distal-effects, there are available strategies to further improve the displayed generalizability of the model. The strategies are: (a) Binary-vectorizing the input without overlap between the channels; (b) Incorporating angular information into GCN; (c) Implementation on high density datasets; (d) Appending error modulating layers downstream; and (e) Incorporating allele information to generalize the VSEPRnet to a pan-specific model. It is also worthy of noting that because the structures of the functional groups are encoded in VSEPRnet, this is applicable to lipids, peptidoglycans, polynucleotides, small molecules, sugars, etc. As long as the size of the molecule is within the limits of the training set, peptide data may be used to train the model while using a small set of peptidoglycans as the test set, for example.

The data and scripts for all the above steps including model building and training are available on GitHub (https://github.com/Sarikaya-Lab-GEMSEC).

## Acknowledgments

We acknowledge the guidance of Kevin Jamieson (Paul G Allen School of Computer Science and Engineering), Marina Meila (Statistics), René Overney (Chemical Engineering and Molecular Engineering and Science Institute) and Deniz T. Yucesoy, (at GEMSEC), all at the University of Washington.

## Supporting information

**S1 Appendix. Description of Channel Inputs to VSEPRnet.** This section describes the information obtained from VSEPR structures of peptides that is sent through each of the 5 channels into the neural network.

**S1 Fig. 5-fold CV data across all alleles.** The 5-fold CV of training set peptides for 9 and 10-Amino Acid long sequences, and their means and standard deviations are tabulated for DeepMHC and VSEPRnet.

**S2 Appendix. Model Comparison of dependency of 5-fold CV on available training data.** This section describes the dependency of 5-fold CV’s obtained from the VSEPRnet and DeepMHC models on available training data.

**S2 Fig. PC, SRCC and AUC metrics from test set.** The Pearson Correlations, Spearman Rank Correlation Coefficients, and Area Under the Curve of test peptides for 9 and 10-Amino Acid long sequences, for DeepMHC and VSEPRnet implementations and their means and standard deviations are tabulated.

## Notes

http://tools.iedb.org/main/datasets

https://github.com/Sarikaya-Lab-GEMSEC

