## Supplementary material for "VSEPRnet: Physical structure encoding of sequence-based biomolecules for functionality prediction: Case study with peptides": S1 Appendix

**S1 Appendix. Description of Channel Inputs to VSEPRnet.** This section describes the information obtained from VSEPR structures of peptides that is sent through each of the 5 channels into the neural network.

Any VSEPR structure is at least parameterized by the Central-Atom (CA), the immediate neighbors bonded to the CA, the length of those bonds, the number of lone-pairs on the CA and the angles made by these bonded neighbors at the central atom. First, protein data bank (PDB) files for each of the naturally occurring Amino Acids (AA) are obtained. These PDB files contain information for each atom in a given AA, including atom type (in terms of atomic number) and cartesian coordinates of the given atom in space. The ProDy library in python is used to traverse through the PDB files and calculate which atoms are bonded to which other atoms using the “findNeighbors” function. Iterating through each of the neighbors of a CA, the bond type of each neighbor bonded to the central atom is obtained, based on prior knowledge of the AA structures. Euclidean distance between one atom and another is calculated by using the well-known three-dimensional distance-formula. After determining the bonds and the bond types, the number of lone pairs on any given CA is inferred based on the number of bonds and the bond-types the given atom has, and the number of valence electrons permissible. Since the Cartesian coordinates of the CA and immediate Bonded-Atoms are known, the dot-product of vectors in 3D space is used to obtain the angle subtended at the CA by any pair of Bonded-Atoms. This is done parallelly for all CA and all combinations of Bonded-Atoms pairs per CA.

Using all of the above information a matrix for a given CA is constructed where each row is a different channel of data, input separately to the neural network for training without overlap. These rows are: a) The first row contains the atomic number of the CA with an index placed after the decimal as a symbolic representation of connectivity. In the tenths place is the index of the residue location and the hundredths place value is the repetition within the residue. For example, the α-Carbon in the 1^st^ AA at the N-terminus is given a value 6.01, whereas the carbon at the center of the planar carboxyl group bonded to the amine group of the 2^nd^ AA, is given a value of 6.00. similarly, the Nitrogen atom at the center of the Amine group of the 1^st^ AA is given a value of 7.00 while that in the 2^nd^ AA is given a value of 7.10. Also, in the first row, to the right of the atomic number with symbolic decimal place-values, there is an integer that represents the number of remaining lone pairs on the CA; b) The second channel contains a list of bond types in order of iteration from N-terminus to C-terminus, traversing side-chains upon encountering an alpha carbon. For example, if the first number in this row is 2, then 1, followed by a 1, then the first bond the computer iterated through was a double bond, second and third are single bonds, for the CA in question. This row contains only integers; c) The third row contains the list of all atoms bonded directly to the CA, in terms of their atomic numbers and symbolic locations, in the same format as the first row; d) The fourth row consists of the corresponding bond lengths for the Bonded-Atoms in the third row, and e) The fifth row contains the bond angles (in degrees) between each pair of immediate neighbors to the CA. Each angle value is subtended at the CA, with pairs of immediate neighbors being the other two vertices. All the five rows are concatenated to make a node-matrix for the CA in question. The largest AA in terms of number of central-atoms is tryptophan, and the largest peptide in the 9-AA or 10-AA case would be a 9 or 10 repeat Tryptophan. Considering this as the baseline, each row in each node-matrix is zero-padded to bring it to a fixed size.

This process is then repeated by treating each non-hydrogen atom as a CA in a peptide with a given sequence, taking into the account the bonds formed between Carboxyl and Amino groups between prior and posterior AA. Each subsequent node matrix is concatenated to the bottom of the previous one, increasing the index (at tenths and hundredths place after decimal in the first and third rows of each node matrix) of each repeating atom type to preserve uniqueness within the whole peptide as well as within the residue. This results in one feature-map per peptide.

While in the training loop, the feature-map is split into 5 channels, with all the first rows per node matrix per feature-map in the first, second rows in second, third rows in the third, fourth rows in the fourth and fifth rows in the fifth. The channels are appended as 1-D vectors one after the other to generate an input tensor that is sent to the neural network as input.
