## Supplementary material for "VSEPRnet: Physical structure encoding of sequence-based biomolecules for functionality prediction: Case study with peptides": S1 Fig

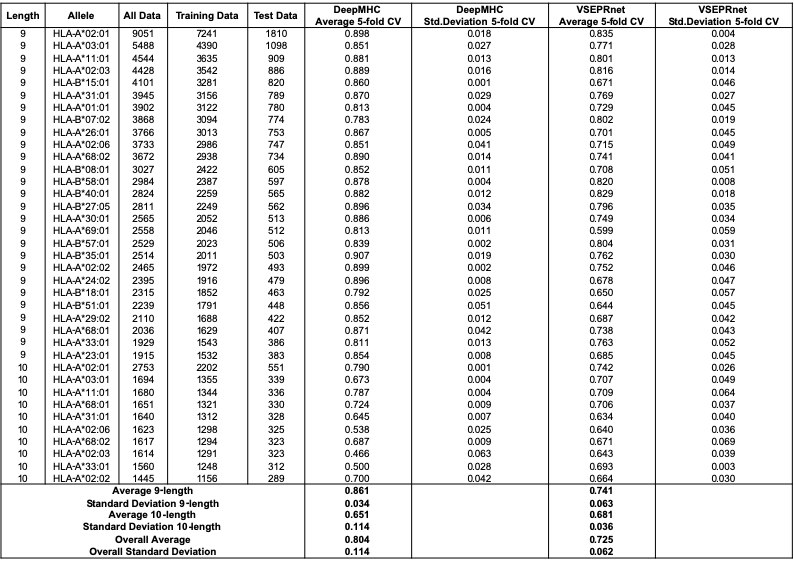

**S1 Fig. 5-fold CV data across all alleles.** The 5-fold CV of training set peptides for 9 and 10-Amino Acid long sequences, and their means and standard deviations are tabulated for DeepMHC and VSEPRnet.
