## Supplementary material for "VSEPRnet: Physical structure encoding of sequence-based biomolecules for functionality prediction: Case study with peptides": S2 Appendix

**S2 Appendix. Model Comparison of dependency of 5-fold CV on available training data.** This section describes the dependency of 5-fold CV’s obtained from the VSEPRnet and DeepMHC models on available training data.

One of the reasons for a marked fall in 5-fold cross validation (CV) for 10-Amibo-Acid (AA) sequences, as outlined in the case of DeepMHC, is a dependency of the model on distal effects arising during convolution which dominate as sequence lengths increase. Upon arranging the alleles in ascending order by number of data-points available to train (S1 Fig) the model, DeepMHC shows a slope of 5.8e^-6^ for 9-AA long peptides, and VSEPRnet shows a slope of 1.15e^-5^, about 1.96 times higher for the same 9-AA long peptides. In the case of 10-AA long peptides, the dependency of 5-fold CV on available training data shows a slope of 1.3e^-4^ for DeepMHC and 5.3e^-5^ for VSEPRnet. DeepMHC thus shows a slope that is 2.48 times higher than VSEPRnet. It is concluded that for the DeepMHC implementation, the change in dependency of 5-fold CV on training data points available is 22.7 times higher for 10-AA long peptides than 9-AA long peptides. In contrast, for the VSEPRnet implementation, it is only 4.65 times higher for 10-AA peptides than 9-AA peptides. This in itself says that for the 10-AA long peptides, VSEPRnet’s 5-fold CV dependency on number of available data-points to train is almost 4.88 times smaller than DeepMHC. An interpretation is that in case of DeepMHC, distal effects along with overfitting (there are fewer 10-AA sequences in the dataset than 9-AA sequences) are responsible for the substantial fall in 5-fold CV. While in the VSEPRnet case, distal effects are either not present due to the implementation of the Efficient Spatial Pyramid scheme in the convolution block, or they do not dominate. Overfitting is the only candidate most likely to cause the slight drop in 5-fold CV between 9-AA and 10-AA sequences. Further research is necessary with similarly large datasets of a variety of sequence lengths, to substantiate the above interpretation.
