## Supplementary material for "VSEPRnet: Physical structure encoding of sequence-based biomolecules for functionality prediction: Case study with peptides": S2 Fig

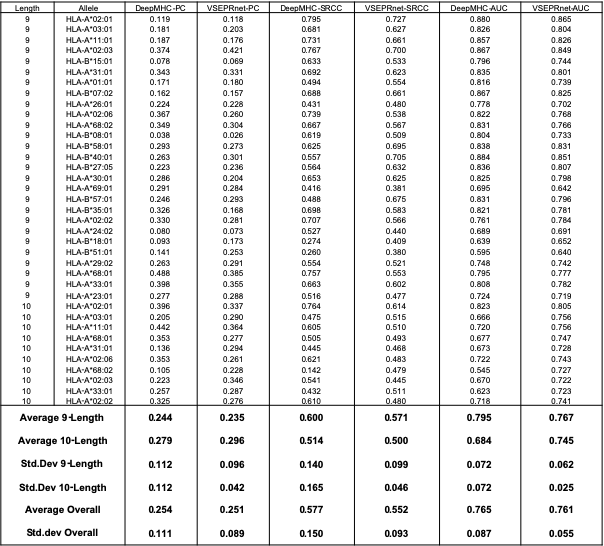

**S2 Fig. PC, SRCC and AUC metrics from test set.** The Pearson Correlations, Spearman Rank Correlation Coefficients, and Area Under the Curve of test peptides for 9 and 10-Amino Acid long sequences, for DeepMHC and VSEPRnet implementations and their means and standard deviations are tabulated.
